# Learning functional conservation between pig and human to decipher evolutionary mechanisms underlying gene expression and complex trait

**DOI:** 10.1101/2023.01.13.523857

**Authors:** Jinghui Li, Tianjing Zhao, Dailu Guan, Zhangyuan Pan, Zhonghao Bai, Jinyan Teng, Zhe Zhang, Zhili Zheng, Jian Zeng, Huaijun Zhou, Lingzhao Fang, Hao Cheng

## Abstract

The assessment of genomic conservation between human and pig at the functional level can help understand and improve the potential of pig as a human biomedical model. To address this, we developed a **<u>Deep</u>** learning-based approach to learn the **<u>G</u>**enomic **<u>C</u>**onservation at the **<u>F</u>**unctional level (DeepGCF) between species by integrating 386 and 374 epigenome and transcriptome profiles from human and pig, respectively. DeepGCF demonstrated a better prediction performance compared to the previous functional conservation prediction method. In addition, we showed that the resulting DeepGCF score captures the functional conservation by examining DeepGCF on chromatin states, sequence ontologies, and regulatory variants. Regions with higher DeepGCF score play a more important role in regulatory activities and show heritability enrichment in human complex traits and diseases. Our DeepGCF approach shows a promising application on the comparison of cross-species functional conservation, and the model framework can be easily adapted to other species. By expanding the model to integrate the functional profiles of multiple species, including human, mouse, pig, cattle, and other livestock animals in the future, the functional conservation information will provide additional insight into the genetic and evolutionary mechanisms behind complex traits and diseases.

## Introduction

Comparative genome not only reveals evolutionary changes at the DNA sequence level^1^, but also helps with the translation of genetic and biological findings across species^2^. Compared to lab organisms like mice, pig is more similar to human in anatomy, physiology, and genome^3^, thus is widely used as a biomedical model for human medicine and genetic diseases, such as drug tests^4^, xenotransplantation^5^, Alzheimer’s disease^6^, breast cancer^7^, and diabetes^8^. To fully recognize the substantial potential of pig as a human biomedical model, it is essential to conduct an extensive comparison of pig and human physiology at the molecular level for assessing to what degree that the genetic and biological findings in pig can be extrapolated to human. Several methods have been proposed to infer the conservation at the DNA sequence level, such as Genomic Evolutionary Rate Profiling (GERP)^9^ and Phylogenetic *P*-values (PhyloP)^10^. However, the conservation at DNA sequence level is not equivalent to the conservation at functional level^11–13^.

The ongoing global efforts on functional annotation of genomes in both humans and livestock, such as the Encyclopedia of DNA Elements^14^, Roadmap Epigenomics projects^15^, the Functional Annotation of Animal Genomes (FAANG)^16^, and Farm animal Genotype-Tissue Expression (FarmGTEx) projects^17^, provide an unprecedented opportunity to quantify the genome conservation across species at the functional level. Previous studies often rely on a single functional profile in one tissue/cell type, such as gene expression^18^ or epigenome^19,20^, to infer the functional conservation of orthologous regions between human and pig. However, integrative analysis of multi-omics is essential for unravelling how biological information encoded in the genome is conserved or diverged across species, as the functional consequence of genomic variants is often modulated at multiple levels of gene regulation across tissues/cells. Artificial neural networks have been applied in the prediction and integration of multi-omics data, such as histone marks, transcription factors, and gene expression, to investigate transcriptional and biochemical impact of DNA sequences and their conservation across species^21,22^. For instance, Kwon and Ernst^22^ developed a neural network model, LECIF, to study human-mouse functional conservation based on multi-omics data from Roadmap and ENCODE databases.

In this study, to systematically evaluate the functional conservation between human and pig, we developed a **<u>Deep</u>** learning-based approach to learn the **<u>G</u>**enomic **<u>C</u>**onservation at the **<u>F</u>**unctional level (DeepGCF) between species. Unlike LECIF using functional genomics data as input, DeepGCF uses both DNA sequences and functional genomics data as input. It thus enables us to predict the impact of sequence mutations on the functional conservation between species. By integrating 386 and 374 epigenome and transcriptome profiles, representing 28 and 21 tissues from human and pig, respectively, DeepGCF captures the conservation of epigenetic features and genes across tissues between human and pig. By further examining expression/splicing quantitative trait loci (e/sQTL) from 54 and 35 tissues in human GTEx^23^ and PigGTEx^24^, respectively, and genome-wide association studies (GWAS) of 80 complex traits/diseases in human, DeepGCF provides novel insights into the evolutionary mechanisms underlying both molecular phenotype and complex trait variation. The DeepGCF model can be easily expanded to multiple species for extensively understanding the genome evolution at functional genomics level when large-scale functional annotation data is available for many other species in the near feature.

## Results

### Overview of the DeepGCF model

In general, the training of DeepGCF model consists of two steps (**Fig. 1**). The first step is to transform the binary functional features to continuous values by training a deep convolutional network implemented in DeepSEA^25^. Binary functional feature is a common data type in the functional genomics filed, which represents whether a genomic base overlapped with functional annotations such as peaks or chromatin states derived from ATAC-Seq and ChIP-Seq. By taking both DNA sequences and binary functional features as inputs, DeepSEA predicts the probabilities of each functional feature at a single-nucleotide resolution. In this study, we collected 309 and 294 genome-wide binary functional annotations from human and pig, respectively (**Supplementary data 1–4**). These represented the chromatin accessibility measured by Assay for Transposase-Accessible Chromatin (ATAC-seq), histone modifications measured by Chromatin Immunoprecipitation sequencing (ChIP-seq) and chromatin states from 26 and 21 tissues in human and pig, respectively. We trained the DeepSEA models and predicted the functional effect of each nucleotide in human and pig separately, which were subsequently used as inputs in the DeepGCF for predicting the functional conservation score between these two species. The performance of DeepSEA was evaluated using an independent validation set and showed a strong predictive power in both species (**Supplementary Fig. 1**).

**Fig. 1.**
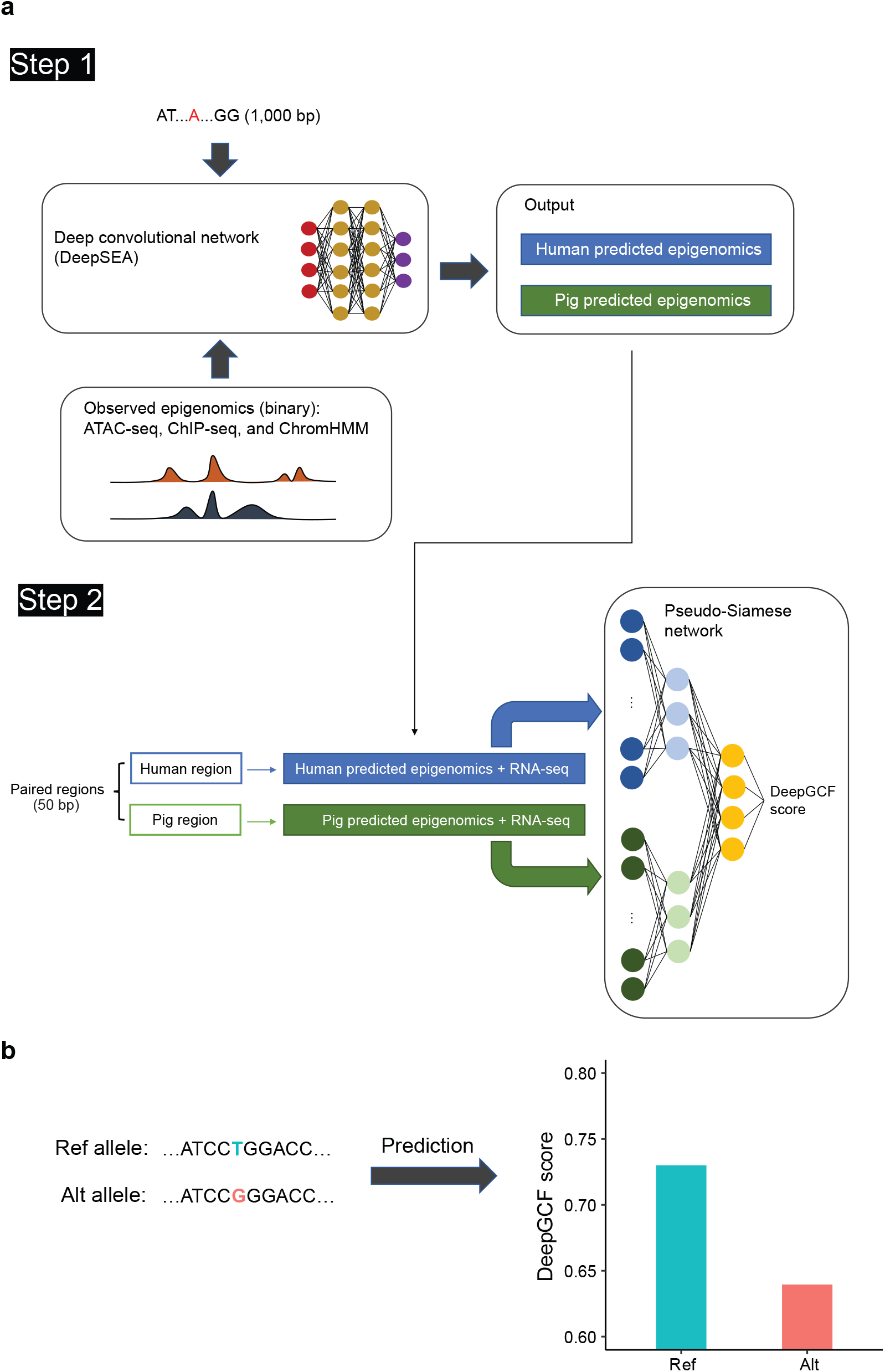
Overview of the DeepGCF model. **a** The learning procedure of DeepGCF model consists of two steps. The first step is to train DeepSEA models in human and pig separately to transform the binary functional features (e.g., peaks called from ATAC-seq and ChIP-seq, and chromatin states predicted from ChromHMM) to continuous values by predicting the functional effects of single nucleotides through centering the target nucleotide at a genomic region of 1,000 bp. The second step is to train a pseudo-Siamese network for predicting whether the paired human-pig regions are orthologous or not using two corresponding vectors of functional effects predicted from DeepSEA and normalized gene expression as inputs. The output, DeepGCF score, is a value between 0 and 1 quantifying the functional conservation of the paired human-pig region. **b** The DeepGCF model can be applied to predict the effect of genome variants on the functional conservation, quantified by changes in DeepGCF scores.

The second step of DeepGCF is to predict the functional conservation score of orthologous regions between human and pig using a supervised deep learning approach, similar to LECIF^22^. We divided the whole-genome alignment between human and pig into non-overlapping 50-bp regions within each alignment block, resulting in 38,961,848 paired alignments (i.e., orthologous regions). We then selected the first base to represent the functional annotation of the 50-bp region, because bases within such a narrow region are likely to have similar functions and the computational burden is greatly lightened by doing so^22^. Apart from the predicted functional effects from DeapSEA, we also included the gene expression values from 77 and 80 RNA-seq datasets as functional annotations, representing 11 and 19 tissues in human and pig, respectively (**Supplementary data 5** and **6**). To train the DeepSEA model, we randomly shifted the human-pig orthologous regions to obtain the same number of non-orthologous pairs. Functional conservation is lack of ground truth, thus as an approximation, we presume that the orthologous regions (coded as 1) are more likely to be functionally conserved than non-orthologous regions (coded as 0). We then trained a pseudo-Siamese neural network model^26^ using both functional effects predicted from DeepSEA and gene expression as inputs (**Fig. 1a**). We weighted non-orthologous regions 50 times more than orthologous ones when training to highlight regions with strong evidence of functional conservation^22^. The output, DeepGCF score, is a value between 0 and 1 quantifying the functional conservation of the paired human-pig region. Furthermore, since the DeepGCF predicts the functional conservation based on the DNA sequence, it allows us to conduct an *in silico* mutagenesis analysis to assess the impact of orthologous variants on the functional conservation between species through investigating the changes of DeepGCF score caused by a mutation (**Fig. 1b**).

### The evaluation of DeepGCF model

The performance of DeepGCF was evaluated by predicting whether the paired human-pig regions of an independent testing set are orthologous or not. Compared to LECIF, which had the areas under receiver operating characteristic curve (AUROC) and precision-recall curve (AUPRC) of 0.80 and 0.79, respectively, DeepGCF showed a better predictive ability with AUROC and AUPRC of 0.89 and 0.87, respectively (**Figs. 2a, b**). Of note, we normalized the gene expression values with a natural logarithm transformation, which showed a better predictive ability than that without a transformation (**Supplementary Fig. 2**). Among all the 38,961,848 orthologous regions between human and pig, only a small percentage (1.2%) exhibited a DeepGCF score greater than 0.8, while more than half with a score less than 0.1 (**Fig. 2c**), consistent with previous findings between human and mice^22^. This result suggests that most of orthologous regions were not functionally conserved between species.

**Fig. 2.**
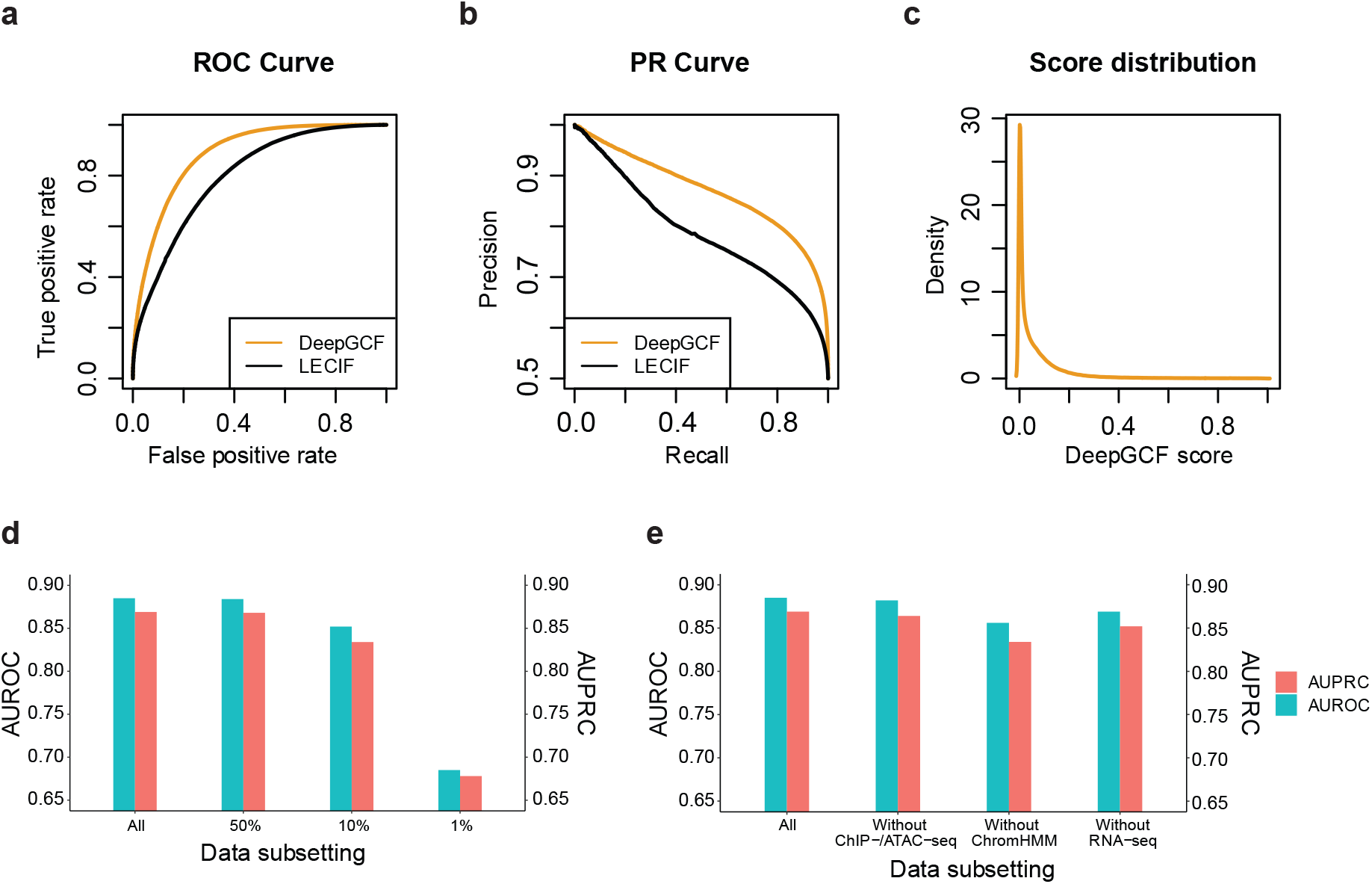
The performance of DeepGCF under different scenarios. **a** Receiver operating characteristic (ROC) curves comparing the performance of DeepGCF (this study) and LECIF^22^ methods. The ROC curve of each method is generated by predicting whether 200,000 pairs randomly selected from the testing set, which included equal number of orthologous and non-orthologous pairs (e.g., randomly mismatched genomics regions), were orthologous or not. **b** Precision-recall (PR) curves generated by similar procedures as the ROC curves. **c** DeepGCF score distribution of all 38,961,848 human-pig orthologues pairs. **d** The areas under receiver operating characteristic curve (AUROC) and precision-recall curve (AUPRC) of DeepGCF using all (Human: 386; Pig: 374), ∼50% (Human: 192; Pig: 187), ∼10% (Human: 52; Pig: 47), and ∼1% (Human: 4; Pig: 4) of human and pig functional features. The subsets of the human and pig features were randomly selected ∼50%, ∼10%, ∼1% from each of ChIP-/ ATAC-seq, ChromHMM, and RNAseq profiles. **e** The AUROC and AUPRC of DeepGCF using all functional features (Human: 386; Pig: 374), features without ChIP-/ATAC-seq (Human: 129; Pig: 84), without ChromHMM (Human: 180; Pig: 210) and without RNA-seq (Human: 77; Pig: 80).

To provide suggestions for researchers who are interested in running the DeepGCF model in other species with limited functional annotation data available, we explored different features that may influence the performance of DeepGCF, including sample size and diversity of functional annotations regarding array and tissue/cell type. When training the model, we downsampled both human and pig functional profiles. We found that using ∼50% (Human: 192; Pig: 187) and ∼10% (Human: 52; Pig: 47) of the functional profiles resulted in similar AUROC (50%: 0.88; 10%: 0.85) and AUPRC (50%: 0.87; 10%: 0.83) values compared to using all the profiles, but using only ∼1% (Human: 4; Pig: 4) of the profiles showed substantially lower AUROC (0.69) and AUPRC (0.68) values (**Fig. 2d**). When leaving one type of functional profiles out, the predictive ability of DeepGCF did not change too much (**Fig. 2e**).

### Relationship between DNA sequence conservation and functional conservation

To fully explore whether DNA sequence conservation indicates functional conservation, we first examined PhyloP scores, which are commonly used to measure the DNA sequence conservation across species^10^. We observed a U-shaped relationship between PhyloP and DeepGCF scores (**Fig. 3a**), demonstrating that both fast-evolving and slow-evolving sequences exhibited a higher functional conservation between species, compared to evolutionary neutral or near-neutral sequences. This agrees with previous findings on comparing individual epigenetic marks and DNA sequence conservation^19,27^. Furthermore, we defined three types of orthologous regions according to their PhyloP and DeepGCF scores to represent the two tails and the bottom of the U curve: 1) regions with both high DeepGCF (> 95^th^ percentile) and PhyloP (> 95^th^ percentile): high D & high P (*n* = 260,281), 2) those with high DeepGCF (> 95^th^ percentile) but low PhyloP (< 5^th^ percentile): high D & low P (*n* = 152,557), and 3) those with low DeepGCF (< 5^th^) and medium PhyloP (between 47.5^th^ and 52.5^th^): low D & med P (*n* = 95,231). By examining sequence classes, which are predicted regulatory activities of DNA sequences in human genome by a deep learning model, Sei, trained on a compendium of 21,907 epigenome profiles^28^, and Gene Ontology (GO) terms, we found that, compared to the whole genome, high D & high P regions were more enriched in promoter, CTCF, and transcription but depleted in enhancer (Binomial test *P* < 0.0001; **Fig. 3b**). Compared to other regions, high D & high P regions showed a higher enrichment in transcription (Binomial test *P* < 0.0001; **Fig. 3b**), and were significantly associated with several RNA-related regulation processes (**Supplementary Data 7**). This indicates the similarities in transcriptional networks between pig and human^18,29^. High D & low P regions were significantly enriched in Polycomb (Binomial test *P* < 0.0001; **Fig. 3b**), in consistency with the fact that some core subunits of Polycomb protein complexes with similar biological functions have shown a weak evolutionary conservation on DNA sequence across species^30^. The low D & med P regions had similar sequence class compositions as the whole genome background except promoter, which was enriched but to a less extent than high D & high P and high D & low P (Binomial test *P* < 0.0001; **Fig. 3b**), and were enriched in fewer GO terms than regions with high DeepGCF (**Supplementary Data 7–9**). In addition, we examined six different sequence ontologies and found that 5’ UTR is the most functionally conserved element, followed by start codon, 3’ UTR, stop codon, exon, and finally intron. This is consistent between both human and pig (**Fig. 3c**).

**Fig. 3.**
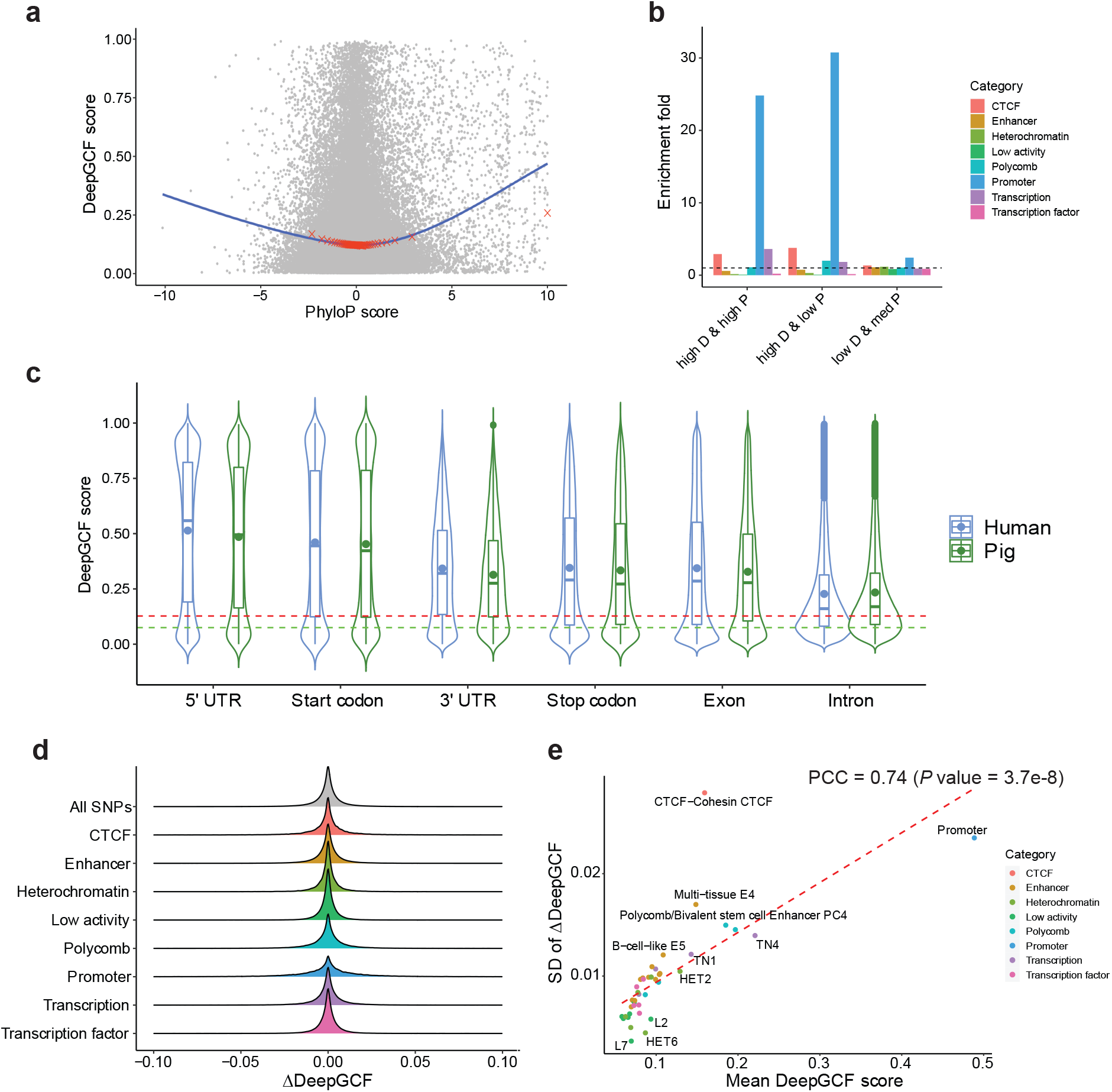
Comparison of functional and sequence conservations. **a** Relationship between DeepGCF score and PhyloP score of 20,000 randomly selected human regions. PhyloP score is based on multiple alignments of 99 vertebrate genomes to the human genome^10^. The blue line is the fitted loess regression and red crosses represents 50 equally-divided percentiles of PhyloP score corresponding to the average of DeepGCF score. **b** Enrichment fold of 8 sequence class categories^28^ for regions with high DeepGCF (> 95^th^ percentile) and high PhyloP (> 95^th^ percentile; high D & high P; *n* = 260,281), regions with high DeepGCF but low PhyloP (< 5^th^ percentile; high D & low P; *n* = 152,557), and regions with low DeepGCF (< 5^th^ percentile) and medium PhyloP (between 47.5^th^ and 52.5^th^ percentile; low D & med P; *n* = 77,848). Enrichment is equal to the proportion of a sequence class category for a type of orthologous regions divided by that for the whole genome. The dashed line (set at 1) represents no enrichment. **c** DeepGCF score distribution of the different sequence ontologies. The red and green dashed lines represent the mean and the median DeepGCF score of the whole genome. The dots inside each box represent the mean DeepGCF score. **d** ΔDeepGCF (DeepGCF after mutation – original DeepGCF) caused by 1,000,000 randomly selected orthologous variants, which are classified into 8 sequence class categories^28^. The red dashed line represents the fitted regression line. **e** The effect of orthologous variants (*n* = 35,575,835) on DeepGCF score of regions in 40 sequence classes^28^, which are classified into 8 categories. The effect was measured by ΔDeepGCF for variants in each sequence class. The SD of ΔDeepGCF for each sequence class quantifies the overall sensitivity of the sequence class to variant effect.

To investigate the impact of orthologous variants on the functional conservation between species, we examined 35,575,835 human SNPs that are located in orthologous regions between human and pig, which were obtained from the 1,000 Genome Project^31^. We used the DeepGCF model trained based on only predicted probabilities of binary features from DeepSEA (i.e., leaving RNA-seq out), as the DeepSEA model does not predict for continuous functional features. The new score predicted from DeepGCF without RNA-seq data had a relatively well agreement with the original DeepGCF score with a Pearson’s correlation coefficient (PCC) of 0.74 (**Supplementary Fig. 3**). To measure the effect of each human SNP on functional conservation, we recomputed the probabilities of binary features for the corresponding orthologous human region due to the SNP mutation and kept the pig probabilities the same, and used the new probabilities to calculate the updated DeepGCF score. The effect on functional conservation is measured by ΔDeepGCF = DeepGCF after SNP mutation – original DeepGCF. By classifying all the orthologous variants into eight categories^28^, we found that most of the variants had a limited effect on the functional conservation (**Fig. 3d**). We further grouped them into 40 sequence classes^28^, and in general, we found that variants in functional features with larger DeepGCF scores showed the stronger effects on the functional conservation between species (**Fig. 3e)**. Promoter and CTCF were more sensitive to variants than other sequence classes. Of note, the average DeepGCF score of CTCF is lower than that of promoter, but it is much more sensitive to genetic mutations regarding the functional conservation, indicating that the genetic disruption of CTCF binding sites (chromatin conformation) may cause strong impacts on functional genome evolution between species by altering the genome topology and consequently the gene expression^32,33^.

### DeepGCF captures the evolutionary characteristics of regulatory elements

To investigate the functional conservation of distinct regulatory elements between pig and human, we first examined the DeepGCF score of 15 chromatin states predicted from 14 pig tissues and 12 human tissues using ChromHMM^19^. We found that strongly active promoters showed the highest DeepGCF scores (i.e., the strongest functional conservation), followed by poised transcription start site (TSS), chromatin states proximal to TSS, enhancers, and finally repressed Polycomb (**Fig. 4a**). This was consistent between human and pig, which agrees with the conservation properties of regulatory elements reported in the previous studies^19,34^. As chromatin states that play important roles in determining the cellular functions may vary among different tissues, we identified strongly active promoters and enhancers that were specific in each of 12 human tissues and 14 pig tissues. Compared to promoters and enhancers shared across all the tissues, tissue-specific ones showed significantly lower DeepGCF scores in both species (Mann–Whitney U test *P* < 2.2e-16), indicative of their faster evolutionary rate (**Fig. 4b**). Among eight common tissues between human and pig, we found that adipose had the strongest functionally conserved promoters in both human and pig, followed by spleen, lung, cortex, liver, and finally stomach (**Supplementary Fig. 4a**). This result suggests pigs could be a good model animal for studying human obesity and metabolic traits^19^. However, the tissue-conservation patterns of enhancers were different from those of promoters and were not consistent between species (**Supplementary Fig. 4b**).

**Fig. 4.**
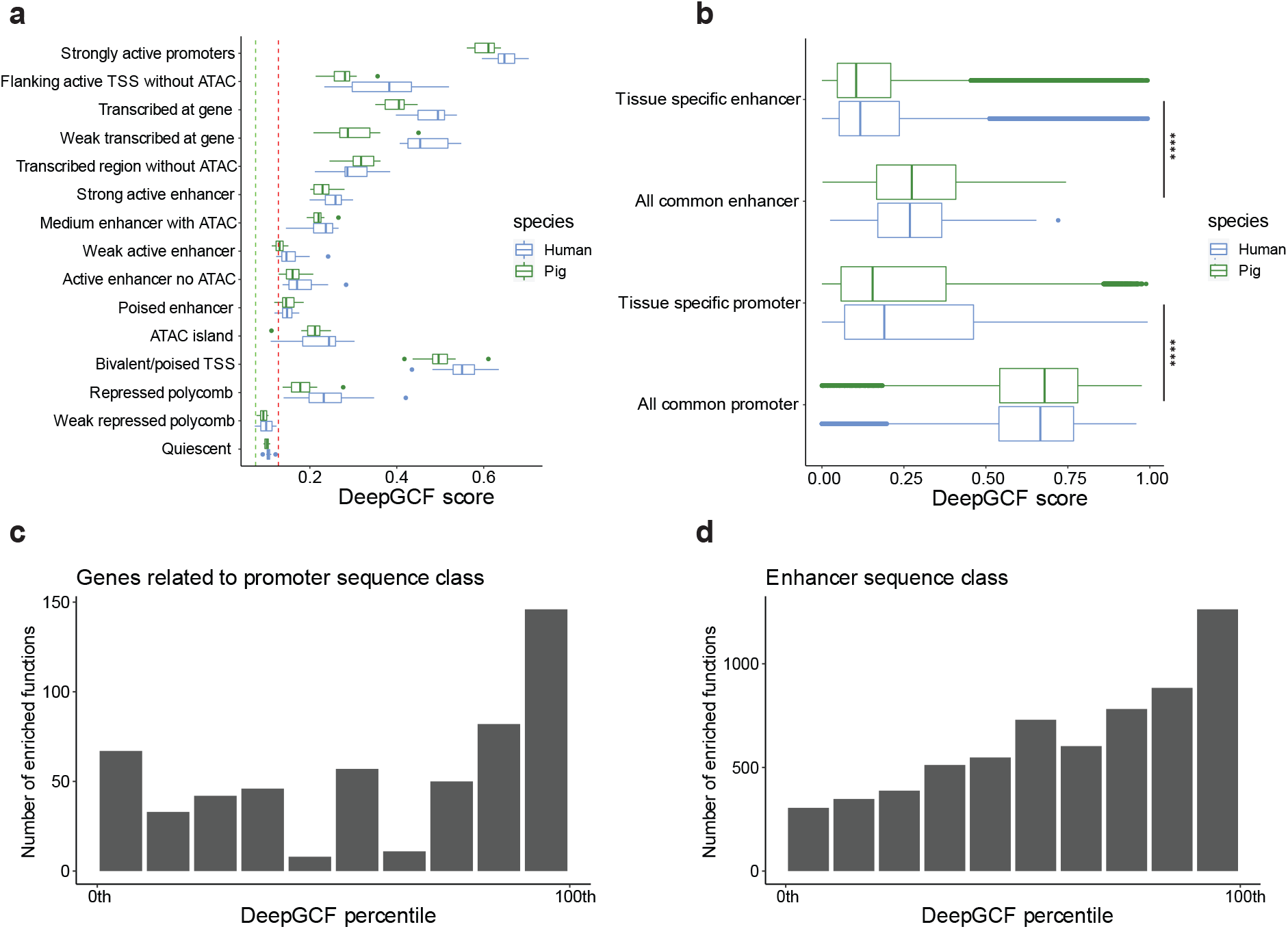
DeepGCF score of genomic regions overlapping with different regulatory elements. **a** Distribution of average DeepGCF scores across human tissues (*n* = 12) and pig tissues (*n* = 14) for each chromatin state. The red and green dashed lines represent the mean and the median DeepGCF score of the whole genome. **b** DeepGCF scores of genomic regions overlapping with tissue-specific strongly active promoter and enhancer for human and pig^19^. “All common” represents promoters/enhancers shared across all tissues. ^****^ denotes Mann–Whitney U test *P* < 2.2e-16. **c** Number of significantly enriched gene ontology terms for human of genes related to promoters annotated by sequence class^28^. The genes were binned by DeepGCF into ten equal-width bins, and the functional enrichment analysis was conducted on each bin. **d** Similar to **c**, except showing the results of enhancers annotated by sequence class^28^.

We further investigated the DeepGCF score on human promoters and enhancers annotated by Sei^28^. We linked a promoter to its potential target gene and then ranked genes with the DeepGCF scores of their promoters (from largest to smallest). We found that top 5% of genes were significantly enriched in basic biological processes, such as anatomical structure development and organ morphogenesis, whereas bottom 5% of genes were significantly enriched in biosynthetic and metabolic process (**Supplementary data 10 and 11**). In addition, we ranked enhancers according to their own DeepGCF scores and investigated the function of top 5% and bottom 5% enhancers. Unlike promoters, top 5% of enhancers exhibited the most significant enrichment in metabolic processes, while bottom 5% of enhancers were significantly enriched in organ growth and development (**Supplementary data 12 and 13**). In general, we found that promoters and enhancers with a higher DeepGCF score were enriched in much more biological processes compared to those with a lower DeepGCF score (**Fig. 4c, d**), which indicates that functionally conserved regions between species tend to be the hotspot of regulatory activities.

### DeepGCF provide insight into the functional conservation of regulatory variants

To explore the functional conservation of regulatory variants, we systematically examined expression QTLs (eQTLs) and splicing QTLs (sQTLs) falling in the orthologous regions in 54 human tissues and 35 pig tissues, respectively. In general, DeepGCF scores of eQTLs and sQTLs were significantly (Mann–Whitney U test *P* < 2.2e-16) higher than the genome background across all the tissues in both human and pig (**Fig. 5a; Supplementary Figs. 5 and 6**), which suggests that regulatory variants are functionally conserved between species^35,36^. Of note, sQTLs showed a higher DeepGCF score than eQTLs in both species (Mann–Whitney U test *P* < 1e-8), probably due to their larger impacts on the transcriptome function (underlying a stronger purifying selection). This is consistent with previous findings that sQTLs were more likely to be enriched in 5’UTR than eQTLs (GTEx, 2020), and 5’ UTR is the most functionally conserved genomic features (**Fig. 2c**). We further observed that eGenes associated with eQTLs having a larger absolute effect on the gene expression had a lower DeepGCF score in both species (**Fig. 5b**), which suggests that orthologous regions with smaller regulatory effects are more likely to be functionally conserved between species, probably due to the stronger purifying selection underlying them^37^. Moreover, regulatory variants influencing more tissues showed higher DeepGCF scores (i.e., more functionally conserved), consistent in human and pig (**Fig. 5c, d**). In addition, the tissue-sharing pattern of orthologous eGenes (PCC = 0.38, *P* value < 2.2e-16) and sGenes (PCC = 0.45, *P* value < 2.2e-16) were positively correlated between human and pig. Altogether, these results indicate that regulatory variants controlling transcriptome function in more tissues tend to be more functionally conserved between species.

**Fig. 5.**
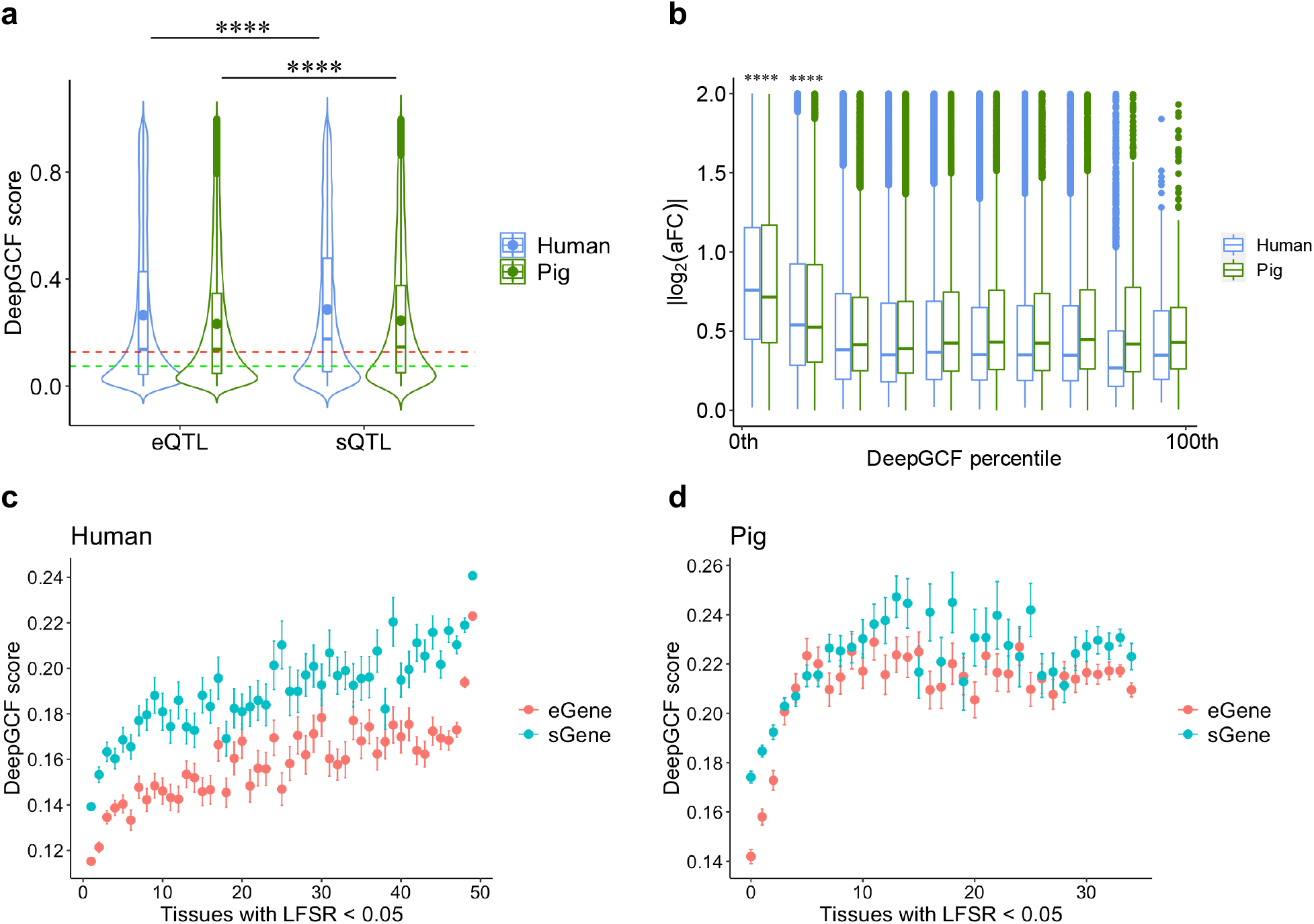
Relationship of DeepGCF score to genetic variants. **a** The distribution of DeepGCF score of eQTLs and sQTLs. The red and green dashed lines represent the mean and the median DeepGCF score of the whole genome. The dots inside each box represent the mean DeepGCF score. ^****^ denotes *P* value < 1e-8 based on a two-sided Mann–Whitney U test. **b** Relationship between the absolute value of eQTL effect size (|log2(aFC)|) and DeepGCF score for eGenes. The genes were binned by DeepGCF into ten equal-width bins for human and pig, respectively. ^****^ denotes the group is different from all other groups with *P* value < 1e-8 based on a Tukey multiple comparison. **c** DeepGCF scores of tissue-sharing e/sGenes from human at local false sign rate (LFSR) < 5% obtained by MashR^50^. **d** Similar to **c**, except showing the results of pig.

We then investigated the DeepGCF scores of 105,461 pathological and likely pathological SNPs obtained from the ClinVar database^38^. A total 98.6% of these SNPs were in the human-pig orthologous regions, consistent with a previous finding that reported more than 98% of pathological variants of Mendelian diseases located in human-mouse orthologous regions^39^. Compared to random orthologous regions, these pathological SNPs were significantly more functionally conserved (Mann–Whitney U test *P* < 2.2e-16; **Fig. 6a**). Like orthologous SNP, we classified the ClinVar SNP into eight sequence class categories^28^ and conducted an *in silico* mutagenesis analysis to predict their impact on the functional conservation. Overall, the average magnitude of variant effect (measured by |ΔDeepGCF|) for pathological and likely pathological mutations is 1.5 times larger than that for random orthologous SNPs (0.0088 versus 0.0058, Mann–Whitney U test *P* < 2.2e-16). In most of cases, the DeepGCF score did not change much after genetic mutations, but the variance of ΔDeepGCF showed a bell-shaped curve regarding the original DeepGCF score, indicating that SNPs with a medium-high DeepGCF (50^th^ to 80^th^ percentile) were more sensitive to pathological mutations than those with lower or higher DeepGCF (**Fig. 6b**). This suggests that the most functionally conserved regions (> 90^th^ percentile) are more tolerable of mutations than less conserved ones (50^th^ to 80^th^ percentile). Most of the ClinVar SNPs were classified as transcription (51.2%), followed by enhancer (16.4%), Polycomb (14.8%), promoter (8.8%), transcription factor (3.3%), and CTCF (2.2%; **Fig. 6c**). Among the ClinVar SNPs with top 5% of |ΔDeepGCF| (> 0.03), there were more SNPs relevant to a decreased DeepGCF (54.4%) than an increased one (45.6%). Moreover, 9 out of 10 ClinVar SNPs with the largest effect on DeepGCF were relevant to a decreased DeepGCF (**Fig. 6c**). In summary, pathological and likely pathological SNPs are located in functionally more conserved regions, and their impact on functional conservation tends to be related to a decreased functional conservation between human and pig.

**Fig. 6.**
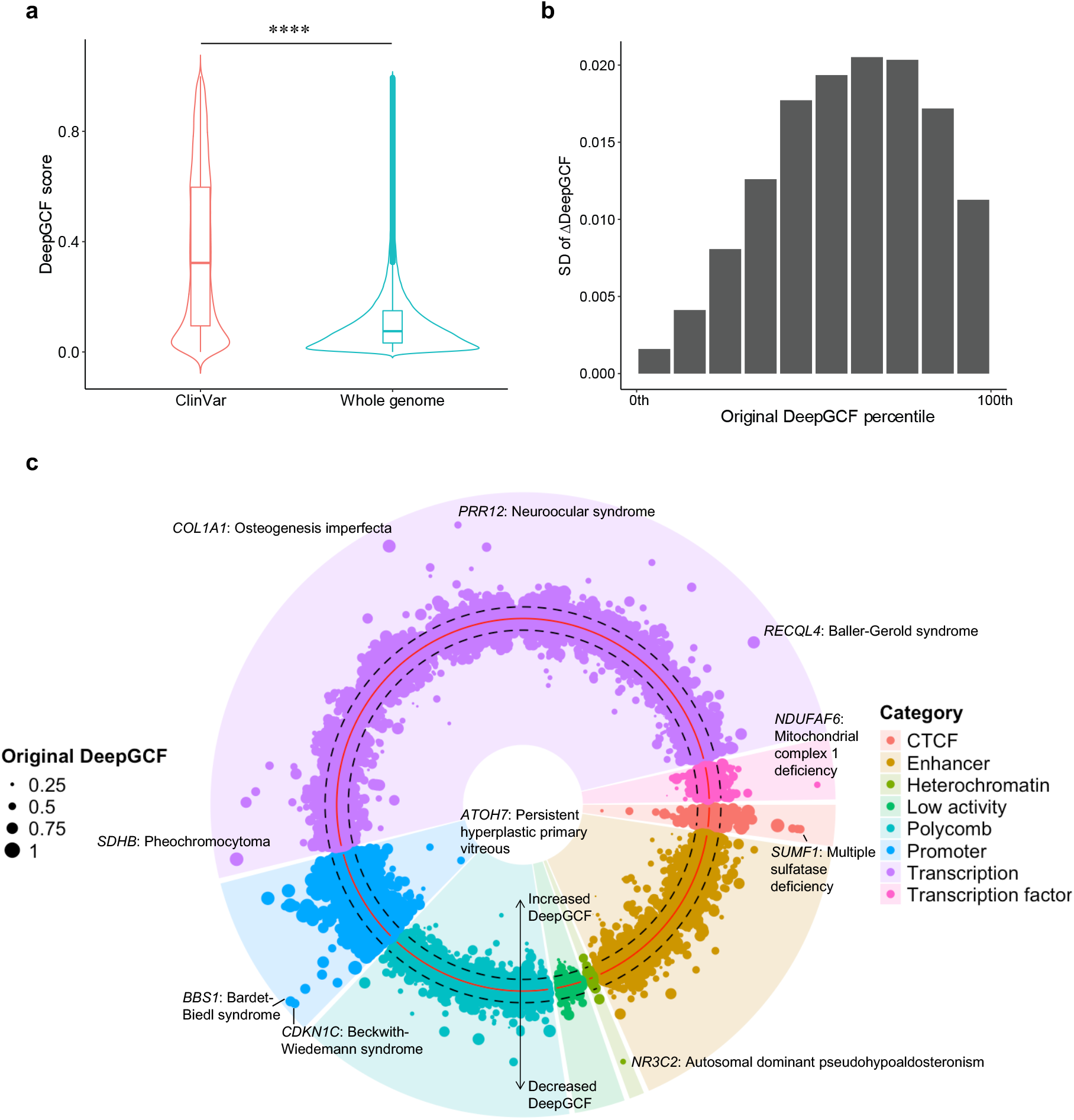
Relationship of conservation score to pathogenic variants. **a** The distribution of DeepGCF scores in pathogenic and likely pathogenic SNPs (*n* = 104,033) obtained from ClinVar^38^, compared to the DeepGCF distribution across the whole genome. ^****^ denotes Mann– Whitney U test *P* < 2.2e-16. **b** SD of ΔDeepGCF (DeepGCF after mutation – original DeepGCF) caused by ClinVar SNPs. The SNPs were binned by their original DeepGCF into ten equal-width bins. **c** ClinVar SNPs classified by sequence class^28^. A polar coordinate system was used, where the radial coordinate indicates the SNP effect on DeepGCF. The red solid circle represents zero DeepGCF change, and two dashed circles represent ± 0.03 of DeepGCF encompassing 95% of SNPs. Each dot represents a SNP and SNPs inside the red circle were predicted to have positive effects (increased DeepGCF), while SNPs outside the red circle were predicted to have negative effects (decreased DeepGCF). Dot size indicates the original DeepGCF. Within each sequence class, SNPs were ordered by chromosomal coordinates. Top 10 SNPs with large impact on DeepGCF associated disease and gene names were annotated.

### Application of DeepGCF on gene mapping and prediction for human complex traits

To investigate whether DeepGCF scores could advance our understanding of the evolutionary basis of complex traits/diseases in human, we conducted a heritability partitioning analysis used the functionally conserved genomic regions (top 5% DeepGCF scores) as a functional annotation, along with 97 existing annotations from the baseline model of LDSC^40,41^, to analyze the GWAS summary statistics from 80 human complex traits/diseases (**Supplementary Data 14**). We found that regions with higher DeepGCF scores explained more heritability of complex traits/diseases (**Fig. 7a**). The heritability of eight complex traits was significantly enriched in functionally conserved regions, with the most enrichment found for coxarthrosis (enrichment = 3.5, FDR = 0.032), followed by varicose veins, height, hypertension, primary hypertension, waist-hip ratio, weight, and BMI (**Supplementary Data 15; Fig. 7b**). Furthermore, we took these eight traits as examples to explore whether DeepCGF can help us with fine-mapping of causal variants. By using functionally conserved regions (top 5% of DeepCGF) as a biological prior in the PolyFun + SuSiE model^42^, we detected 33, 22, and 17 additional putative causal variants (PIP > 0.95 and *P* < 5e-8) compared to the SuSiE model only without any priors in height, BMI and weight, respectively (**Fig. 7c**, Supplementary Data 16). We further incorporated DeepCGF in SBayesRC^43^ model to conduct polygenic score prediction for 20 human complex traits (**Supplementary Data 17**). On average, the relative prediction accuracy increased by 0.56% (**Fig. 7d; Supplementary Data 18**), and the largest increase was observed on waist-hip ratio (3.5%), followed by body weight (1.7%). Altogether, our results showed that DeepGCF provide additional insights into the genetic and evolutionary basis of complex phenotypes.

**Fig. 7.**
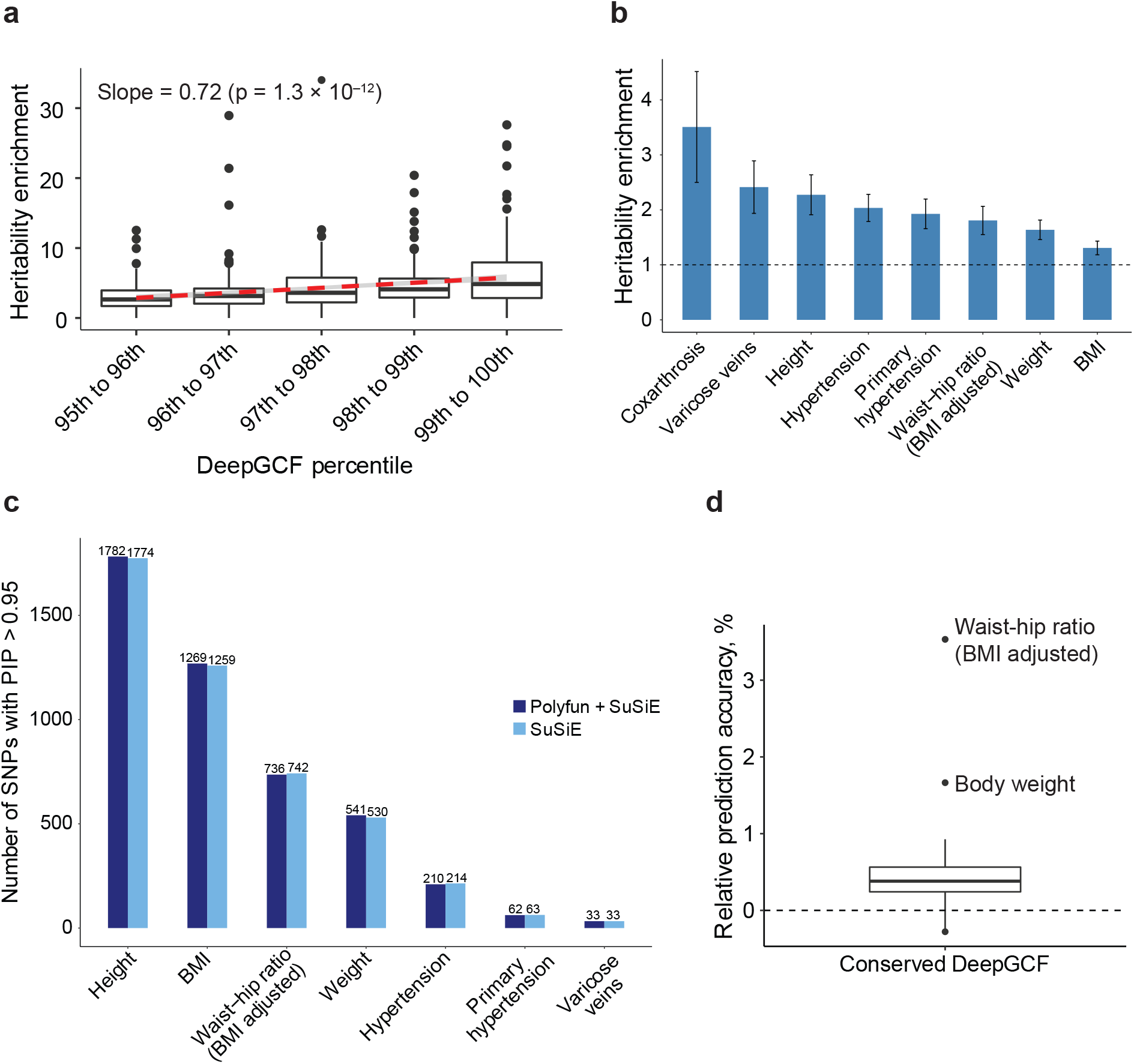
Application of DeepGCF on complex traits/diseases in human. **a** Heritability enrichment calculated by LDSC for 80 human traits using functionally conserved regions (top 5% DeepGCF). The regions were divided into 5 equal equal-width bins and the heritability enrichment of all traits was calculated for each bin. The dashed red line is the fitted regression line between heritability enrichment and DeepGCF percentile, and the grey area is the 95% confidence interval. **b** Significant heritability enrichment explained by functionally conserved regions in 8 human traits. **c** The number of putatitive SNPs (PIP > 0.95 and *P* < 5e-8) identified by PolyFun + SuSiE using functionally conserved regions as a prior and SuSiE without priors for 7 human traits. **d** The relative prediction accuracy of PRS for 20 human complex traits using functionally conserved regions as a prior in SBayesRC^43^. Relative prediction accuracy is equal to (prediction accuracy using the prior – prediction accuracy without priors) / prediction accuracy without priors. A relative prediction accuracy > 0 (dashed line) indicates an accuracy higher than without priors.

## Discussion

In this study, we developed a two-step neural network approach, DeepGCF, to evaluate the genomic conservation at the functional level between human and pig. DeepGCF shares a similar model structure as LECIF^22^ in the evaluation of functional conservation by comparing the epigenome and gene expression profiles of orthologous regions between two species. But instead of using binary epigenome profiles as the direct inputs, DeepGCF first predicts their functional effects (i.e., the continuous probability score of each epigenome binary feature) using DeepSEA^25^, and then use them as the input to predict the functional conservation between species. Compared to the LECIF approach, DeepSEA showed a better performance in the ortholog prediction, probably due to a higher resolution of the model input. Similar to LECIF, we found that the performance of DeepGCF was not sensitive to the number of functional features, indicating that DeepGCF could be applied on other species where functional features are not abundant.

We demonstrated that functional conservation is different from sequence conservation. The relationship between DeepGCF and PhyloP scores confirms the U shape relationship between functional and sequence conservation. By examining DeepGCF on chromatin states, sequence ontologies, and regulatory variants, we verified that DeepGCF captures the functional conservation of genome, and regions with higher DeepGCF play a more important role in regulatory activities. We thereby expected DeepGCF to be useful in explaining complex traits and diseases. The heritability enrichment and polygenic prediction accuracy brought by functionally conserved regions were limited, this may because we only considered functional conservation between human and pig compared to sequence conservation which were obtained based on over 100 species^44^. With the increasing amount of epigenome and gene expression data in other species in the near future, we could identify the core functionally conserved regions by expanding the DeepGCF model structure to integrate functional profiles from multiple species. Another limitation is that the functional conservation of the same sequence segment in different tissues and cell types should be conceptually different, which could not be distinguished by the current DeepGCF score. One ideal way to obtain tissue- and cell-type-specific DeepGCF scores is to train a different model on each tissue and cell type using the respective data. However, the current volume of functional profiles, particularly in pig, does not support the development of tissue- and cell-type-specific DeepGCF models.

Despite the limitations, the DeepGCF approach shows a promising application on the comparison of cross-species functional conservation. The model framework can be easily adapted to other species. Our future work will focus on expanding the model to the comparison of multiple species, including human, mouse, pig, cattle, and other livestock animals. The functional conservation information among different species will provide additional insight into the genetic and evolutionary mechanisms behind complex traits and diseases, analogous to the sequence conservation among vertebrate animals provided by such as PhyloP score.

## Methods

### Genome alignment

We used the chained and netted alignments of human (GRCh38) and pig (susScr11) genome assemblies from the UCSC genome browser^45^. The assemblies were aligned by the lastz alignment program^46^ using human as the reference.

### Model inputs

We divided the whole-genome alignment between human and pig into non-overlapping 50-bp regions within each alignment block, resulting in 38,961,848 orthologous pairs. If an alignment block ended shorter than a 50-bp window, the window was truncated to the end of the block, which resulted in some regions smaller than 50 bp. For each orthologous pair, we collected the corresponding functional features, including chromatin accessibility measured by Assay for Transposase-Accessible Chromatin (ATAC-seq), histone modifications measured by Chromatin Immunoprecipitation sequencing (ChIP-seq), chromatin state annotations (ChromHMM), and gene expression measured by RNA-seq for human and pig from public resources, including ENCODE^14^ and public literatures^19,20^. We only collected the functional data at the tissue level for human, and merged those of the same data type from the same tissue, so that the total number of human features were close to pig. For human, there were 604 ChIP-seq and ATAC-seq files merged into 129 features, 12 ChromHMM files of 15 chromatin states (12 × 15 = 180 features), and 77 RNA-seq features, which resulted in 386 functional annotations. For pig, there were 287 ChIP-seq and ATAC-seq files merged into 84 features, 14 ChromHMM files of 15 chromatin states (14 × 15 = 210 features), and 80 RNA-seq features, which resulted in 374 functional annotations. Details of features from each data type are reported in Supplementary Data 1–6.

### Prediction of binary functional features based on DeepSEA

We trained two DeepSEA models to predict the binary functional features, including ATAC-seq, ChIP-seq and chromatin state annotations, of human and pig using the PyTorch-based package, Selene^47^. We used the peak calls of ATAC-seq and ChIP-seq, and one-hot encoded chromatin state annotations as the training input. We then trained the model based on a sequence region of 1,000 bp, and the feature must take up 50% of the center bin (200 bp) for it to be considered a feature annotated to that sequence. All the hyperparameters were set as default (Supplementary Data 19). We created a validation set using the data from chromosomes 6 and 7 for early stopping during training, a test set using the data from chromosomes 8 and 9 for the generation of the receiver operating characteristic (ROC) and precision-recall (PR) curves, and a training set using the rest data. We then predicted the probability of each binary feature using the trained model for the first base of all the paired regions that were at most 50 bp.

### Data subsets for training and evaluation

We divided the entire data into the training, validation, and prediction sets based on the chromosome number. To predict the DeepGCF score of human regions from even and X chromosomes (prediction set), and the corresponding paired pig regions, we trained a DeepGCF model based on paired regions from a subset of odd chromosomes of human and pig. We created a validation set also from another subset of odd chromosomes (not overlapping with the training set) for the hyper-parameter tuning and early stopping during training. We used a subset of the test set to generate the ROC and PR curves. To predict the DeepGCF score of human regions from odd chromosomes and the corresponding paired pig regions, we created training and validation set similarly as above, except from even chromosomes. We excluded Y and mitochondrial chromosomes in this study. Detailed division of each set is shown in Supplementary Data 20.

### DeepGCF training

Before training the DeepGCF model, we first randomly paired up the human-pig orthologous regions to get the same number of non-orthologous pairs in the training set. We then trained the DeepGCF model with a pseudo-Siamese architecture as the LECIF model^22^. In our pseudo-Siamese neural network, for each orthologous/non-orthologous pair, two input vectors containing the human and pig binary features (probabilities between 0 and 1) predicted from DeepSEA and normalized RNAseq data (also between 0 and 1) were connected to the human and pig subnetworks, respectively (Fig. 1). We performed a natural logarithm transformation on RNAseq data given the large range before normalizing. The two subnetworks were then fully connected to a final subnetwork, which generated the output prediction. We weighted non-orthologous pairs 50 times more than orthologous ones during the training process.

We conducted a random grid search for hyper-parameters, including number of layers in each subnetwork and the final subnetwork, number of neurons in each layer, learning rate, batch size, and dropout rate. We generated 100 combinations of hyper-parameters randomly selected from the candidate parameter pool (Supplementary Data 21), using each combination to train a DeepGCF model based on the same random subset of 1 million aligned and 1 million unaligned human-pig pairs from the training set. We then selected the combination of hyper-parameters that maximized the AUROC on the validation set to train the final model based on the whole training set. We stopped training if there was no improvement in AUROC over three epochs on the validation set for both hyper-parameter search and training, otherwise the training stopped when reaching the maximum number of epochs, which was set to be 100.

### Human-pig orthologous SNPs

In total 73,257,633 human biallelic SNPs (GRCh38) were obtained from 1,000 Genome Project^31^. Their positions were lifted to corresponding orthologous positions in the pig genomes (SusScr11) using the UCSC liftover utility with chain files available from the UCSC website^45^, which resulted in 35,575,835 orthologous SNPs.

### Function enrichment

To explore the Gene Ontology terms of genomic regions (e.g., enhancers), we used the GREAT tool^48^ with default parameters and a cut-off of FDR < 0.05 for both the binomial and the hypergeometric distribution-based tests.

### Tissue specific chromatin state

For each chromatin state, we first used the merge function of BEDtools^49^ to merge any regulatory regions between two tissues overlapping by at least 1 bp across all tissues. Then for strongly active enhancer and promoter in each tissue, if a region is active in only one tissue and does not overlap with any active regions in other tissues, we define the region as tissue specific regulatory element. If a region is active in all tissues (i.e., overlaps across all tissues), we define the region as “all common” regulatory element.

### Tissue-sharing of e/sGene

To explore how e/sGenes are shared across all tissues, we performed the meta-analysis of e/sGenes using MashR (v0.2.57)^50^. We used the slope and the standard error of slope of top e/sQTL of genes (missing slopes were set to be 0 with standard error of 1) across 49 tissues from GTEx (v8)^23^ for human and 34 tissues from PigGTEx databases^24^ for pig as the input. We then obtained the estimate of effect size and the corresponding significance (local false sign rate, LFSR) from the mash function. An e/sGene was considered active in a tissue if LFSR < 0.05.

### DeepGCF score for genes

We obtained the gene boundaries of human and pig genes from Ensembl release 107 (GRCh38 for human and Sscrofa11 for pig), and extended them by 35 kb upstream and 10 kb downstream to include probable cis-regulatory regions^51^. We then compute the DeepGCF score for genes based on the average score of all orthologous regions overlapping with the gene and the extended regions. For human genes linked to promoter sequence class, we identified a promoter’s potential target gene if the distance between the promoter and the TSS of a gene is less than 2 kb, yielding a total of 12,044 promoter-gene pairs.

### Heritability partitioning analysis

We collected the GWAS summary statistics of 80 human complex traits from the UK Biobank and public literatures (Supplementary Data 14). We ran the LD-score regression software ldsc (v1.0.1)^41^ to partition the heritability based on two sets of annotations: 1) one binary annotation of functionally conserved regions (top 5% of DeepGCF) and 2) five binary annotations dividing the top 5% DeepGCF into 5 equal-width bins based on percentiles. Both sets of annotations were analyzed with a baseline including 97 annotations^40^. Heritability enrichment was calculated as the proportion of trait heritability contributed by SNPs in the annotation over the proportion of SNPs in that annotation.

### Fine-mapping analysis

We first used PolyFun^42^ to compute SNP prior causal probabilities based on the annotation of functional conservation (top 5% DeepGCF). These prior causal probabilities were then used as priors in SuSiE^52^ for the fine-mapping analysis. To compare fine-mapping using functional conservation as prior with not using it, we also performed a fine-mapping analysis using SuSiE alone, which only took LD information into account. A SNP is identified to be putative causal if the posterior causal probability (PIP) is greater than 0.95 and the *P*-value in GWAS is smaller than 5e-8.

### Polygenic prediction

We incorporated functional conservation as a prior in polygenic prediction using the software SBayesRC^43^. The GWAS summary statistics of 20 complex traits from UK Biobank (Supplementary Data 17) were analyzed using ∼7 million common SNPs with and without one annotation of functional conservation (top 5% DeepGCF). To compare the prediction accuracy, we partitioned the total sample into ten equal-sized disjoint subsamples. For each fold, we retained one subsample as the validation set and other remaining nine subsamples as the training set. We calculated the polygenic score (PGS) using genotypes from an independent validation set in each fold and obtained the prediction R^2^ from linear regression of true phenotype on the PGS. We then calculated the relative prediction accuracy by (R_0_^2^ – RD2) / R_0_^2^, where R_0_^2^ is the prediction R^2^ without any priors, and R_D_^2^ is the prediction R^2^ using functional conservation as a prior.

## Data availability

The DeepGCF score for human-pig orthologous regions are publicly available for download without restrictions from https://github.com/liangend/DeepGCF. All epigenomic and gene expression data used for model training can be found in Supplementary data 1–6. Orthologous SNPs between human and pig are from the 1,000 Genome Project (http://ftp.1000genomes.ebi.ac.uk/vol1/ftp/data_collections/1000_genomes_project/release/20181203_biallelic_SNV). GWAS summary statistics used for LDSC analysis are from UK Biobank (http://www.ukbiobank.ac.uk), with details showing in Supplementary data 14. Summary statistics and genotype used for polygenic score prediction from UK Biobank (http://www.ukbiobank.ac.uk) are available through formal application.

## Code availability

The code of DeepGCF is available at https://github.com/liangend/DeepGCF.

## Notes

### Competing Interest Statement

The authors have declared no competing interest.

### Summary of Updates

Changed some typos.

