## Supplementary Figures for "Learning functional conservation between pig and human to decipher evolutionary mechanisms underlying gene expression and complex trait"

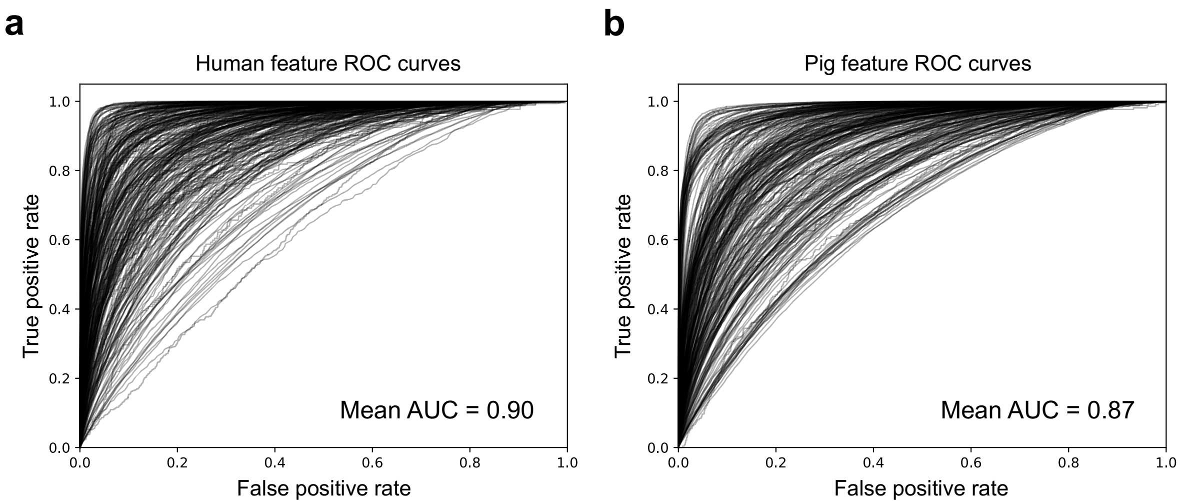


**Supplementary Fig. 1 Prediction accuracy of DeepSEA on binary features.** Receiver operating characteristic (ROC) curve of DeepSEA prediction on **a** human and **b** pig binary features, including ATAC-seq, ChIP-seq and ChromHMM state. The curves were generated by predicting those features on 120,000 samples randomly drawn from the testing set. Each curve represents the prediction accuracy evaluated on one binary feature.

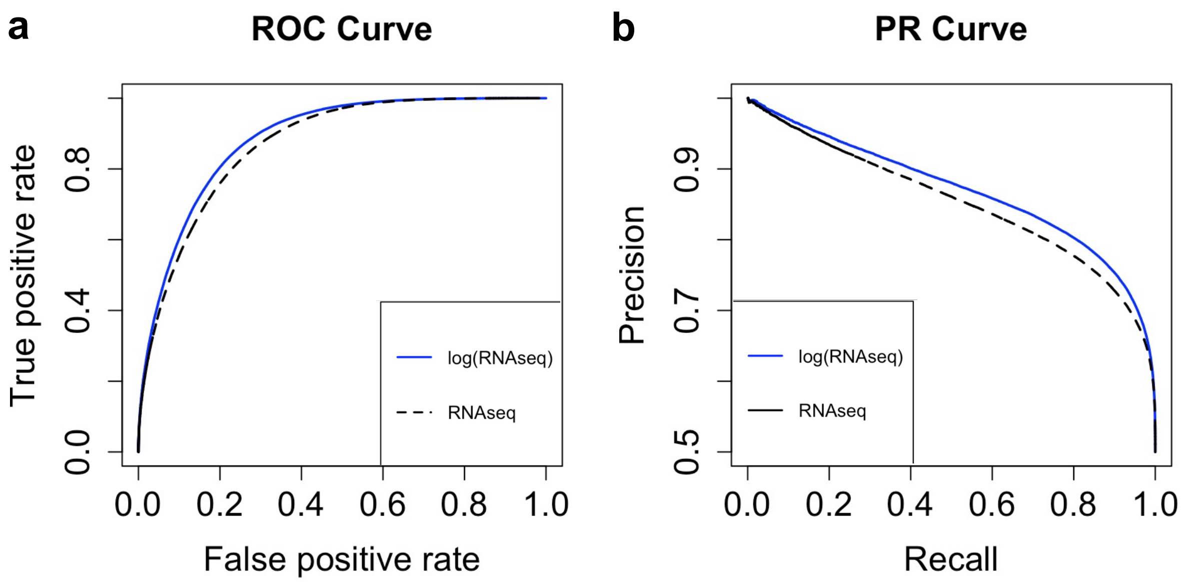


**Supplementary Fig. 2 Comparison of DeepGCF using normalized RNAseq data with or without natural logarithm transformation. a** Receiver operating characteristic (ROC) curve and **b** precision-recall (PR) curve. The curves were generated by predicting the alignment of 200,000 aligned and unaligned pairs, which were randomly drawn from the testing set. The areas under ROC curve and PR curve of DeepGCF using normalized RNAseq data after natural logarithm transformation were 0.89 and 0.87, while those of without natural logarithm transformation were 0.87 and 0.85, respectively.

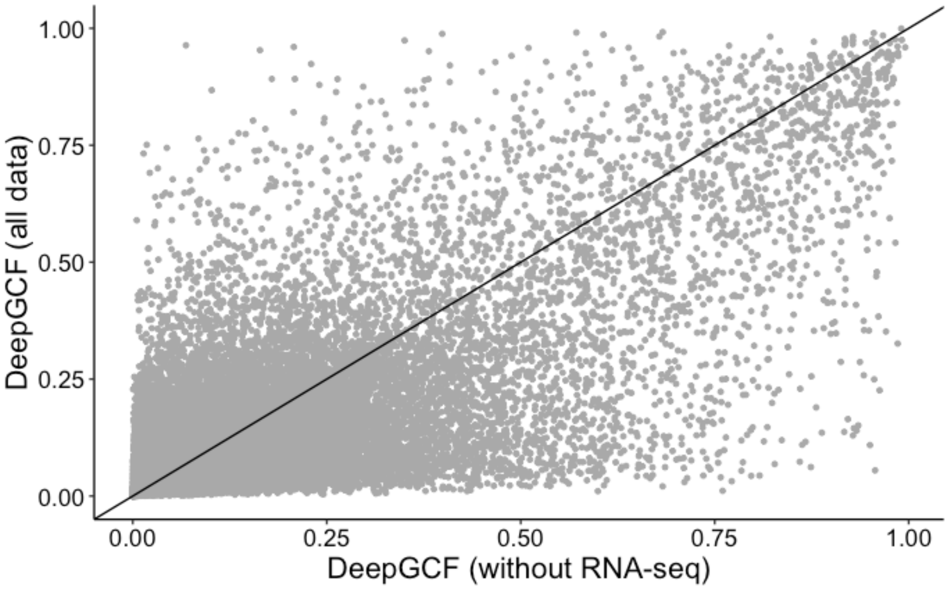


**Supplementary Fig. 3 Comparison between DeepGCF trained with and without RNA-seq data.** The DeepGCF score of 50,000 randomly selected regions predicted using the model trained with/without RNA-seq data of human and pig. The Pearson's correlation coefficient = 0.74.

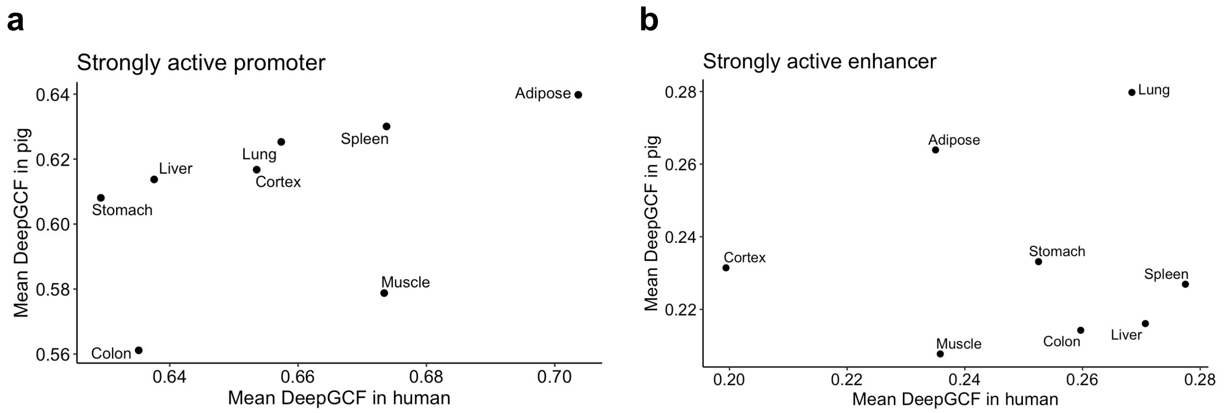


**Supplementary Fig. 4 Mean DeepGCF of strongly active promoters and enhancers. a** Mean DeepGCF scores of genomic regions overlapping with strongly active promoter across 8 common tissues from human and pig. **b** Similar to **c**, except showing the results of strongly active enhancer.

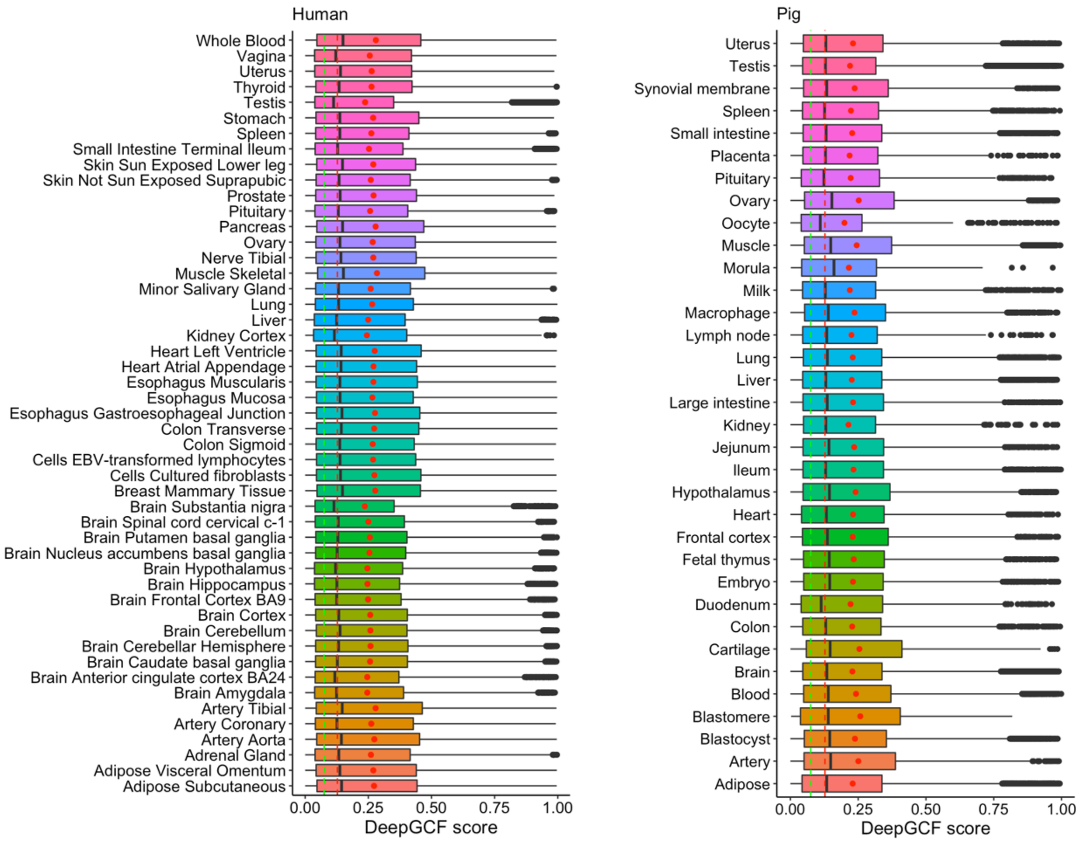


**Supplementary Fig. 5 DeepGCF of eQTLs.** The distribution of DeepGCF score in regions overlapping with eQTLs across each tissue and cell type for human (*n* = 475,829) and pig (*n* = 145,810). Red dots represent the mean DeepGCF score of eQTLs in the corresponding tissue or cell type. The red dashed line represents the average score across the genome, and the green dashed line represents the median.

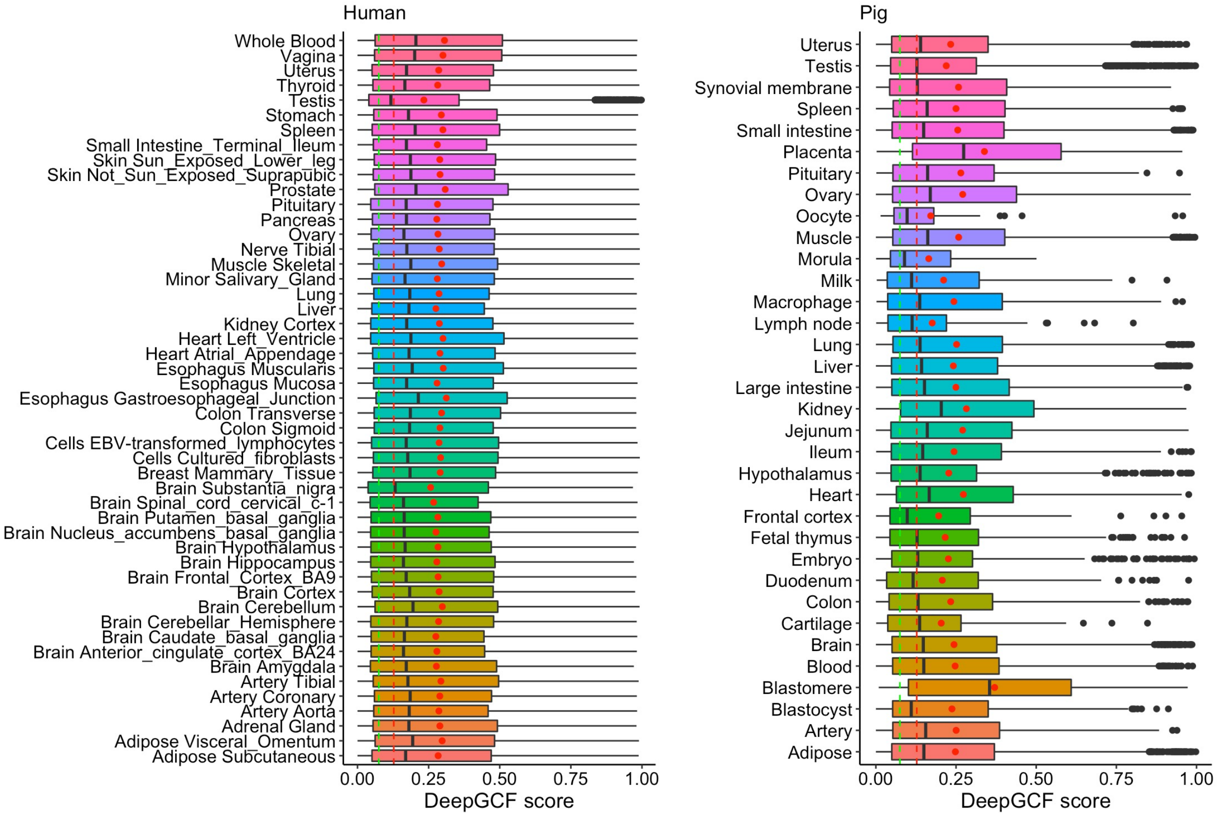


**Supplementary Fig. 6 DeepGCF of sQTLs.** The distribution of DeepGCF score in regions overlapping with sQTLs across each tissue and cell type for human (*n* = 76,047) and pig (*n* = 21,556). Red dots represent the mean DeepGCF score of sQTLs in the corresponding tissue or cell type. The red dashed line represents the average score across the genome, and the green dashed line represents the median.
